# PAVOOC: Designing CRISPR sgRNAs using 3D protein structures and functional domain annotation

**DOI:** 10.1101/398859

**Authors:** Moritz Schaefer, Dr. Djork-Arné Clevert, Dr. Bertram Weiss, Dr. Andreas Steffen

**Affiliations:** Computer Science, TU Berlin, D-10623 Berlin, Germany; Bioinformatics, Bayer AG, D-13342 Berlin, Germany

## Abstract

**Summary:** sgRNAs targeting the same gene can significantly vary in terms of efficacy and specificity. PAVOOC (Prediction And Visualization of On- and Off-targets for CRISPR) is a web-based CRISPR sgRNA design tool that employs state-of-the art machine learning models to prioritize most effective candidate sgRNAs. In contrast to other tools, it maps sgRNAs to functional domains and protein structures and visualizes cut sites on corresponding protein crystal structures. Furthermore, PAVOOC supports HDR template generation for gene editing experiments and the visualization of the mutated amino acids in 3D.

**Availability and Implementation:** PAVOOC is available under https://pavooc.me and accessible using current browsers (Chrome/Chromium recommended). The source code is hosted at github.com/moritzschaefer/pavooc under the *MIT License*. The backend, including data processing steps, and the frontend is implemented in Python 3 and ReactJS respectively. All components run in a simple Docker environment.

## 1 Introduction

The discovery of the CRISPR/Cas system ([8], [2]) was a breakthrough in the area of genome editing. An important application of CRISPR/Cas is to induce a targeted knockout (KO) of a gene of interest. Such KO experiments can help to study the essentiality of the targeted genes in given cellular contexts (e.g. a cancer cell line bearing certain genomic alterations) and ultimately support the validation of a new drug target ([12]). Shi et al. showed, that the effect of a CRISPR based KO can be boosted by targeting functionally relevant regions of a protein, as in these regions in-frame mutations (indels) are more likely to induce a significant effect than in non-functional regions ([14]). Another application of CRISPR/Cas is to precisely introduce missense mutations into a genome and study the resulting effects of the perturbations. In both applications single-guide RNAs (sgRNAs) are used to direct the Cas9 enzyme towards the genomic region of interest, such that the Cas9 can cut the DNA at the targeted position. For the genome editing experiments in addition a template sequence needs to be provided that contains the desired nucleotide sequence.

A number of tools have been published that facilitate and automate the design of sgRNAs for CRISPR KO experiments ([11], [7], [9], [15]). In this application note we present PAVOOC (Prediction And Visualization of On- and Off-targets for CRISPR) – a modern web application to support wet lab biologists in designing and selecting optimal sgRNAs and template sequences for KO and genome editing experiments using machine learning-based on-and off-target scoring, multi-attribute ranking, protein structure mapping of the cut sites and integration of cancer cell line data.

## 2 Materials and methods

PAVOOC is a web application that allows to design and visualize sgRNAs for gene KO and gene editing experiments. For KO experiments, a set of genes has to be provided (in form of symbols or Ensembl identifiers). PAVOOC then generates a table that contains a user-defined number of sgRNAs for each of these genes. These sgRNAs are prioritized based on a scoring function that combines weighted on-and an off-target scores. The on-target score is calculated using the Azimuth model, whereas the Cutting Frequency Determination (CFD) score is used to assess off-target effects ([4]). Optionally, the genomic background of a cancer cell line can be taken into account for the sgRNA prioritization.

It is possible to further analyze and modify the sgRNA selection for a gene in a detail view (see Figure 1). The detail view consists of three synchronized panels: The “LineUp” ranking table on the upper right, the protein structure view on the upper left and the sequence view on the bottom panel of the page. The LineUp ([5]) based sgRNA ranking table allows an individual adjustment of the weights for the on- and off-target scores in order to prioritize the sgRNAs accordingly. For each sgRNA, the LineUp table displays whether the targeted genomic region lies within a protein domain and whether the optionally selected cancer cell line contains a single nucleotide variation (SNV) at that position. The sequence view on the bottom shows the gene annotation, all targeted regions of the sgRNAs, protein domains and cancer cell line alteration data in order to support the tailored sgRNA design for a cell line under study. On the left side, available 3D protein structures are shown and sgRNA-related cut sites are mapped and highlighted on the structure. In this way, the user can assess the position of the Cas9 cut position on the protein structure and thus prioritize sgRNAs that are more likely to affect functionally relevant regions of a gene. Furthermore, when designing gene editing experiments, the structure view enables amino acid editing and displaying the designed alterations directly on the protein structure.

**Figure 1.**
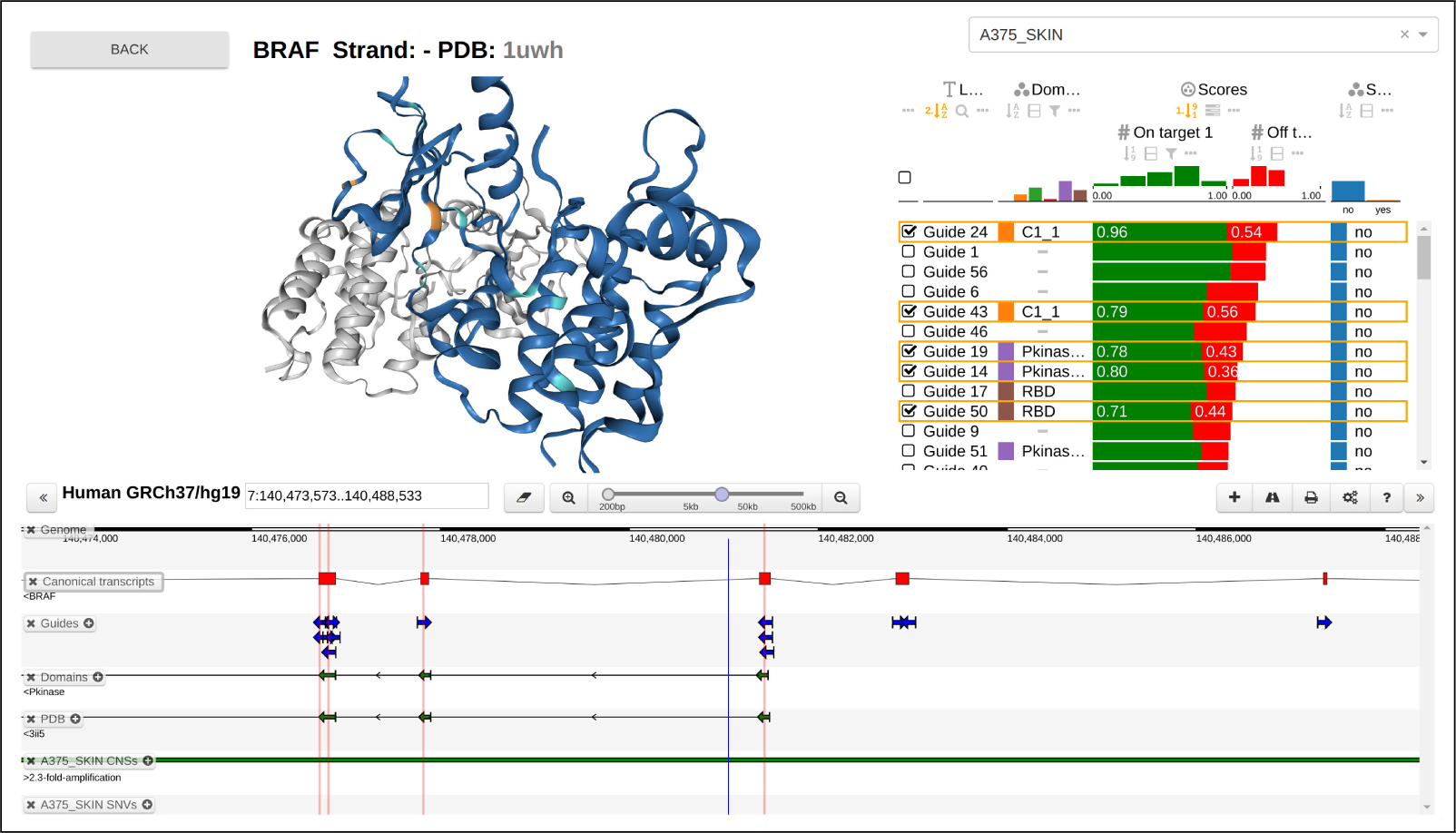
Screenshot of the view to edit sgRNA selection, showing the LineUp ranking table and the protein structure view on the upper panels and the sequence viewer on the bottom panel.

We integrated genomic sequence data from UCSC in version hg19 ([3]). The genomic annotations, including genes, transcripts and exons were taken from the GENCODE project ([6]). Cancer cell line alteration data was taken from the Cancer Cell Line Encyclopedia ([1]) (based on hg19). In order to facilitate the mapping between genomic and protein coordinates we used the canonical transcript from APPRIS ([13]) only. Exons which are not contained in that transcript are not considered in our application. SIFTS ([16]) mappings are used to derive genomic coordinates of PDB structures.

The data shown in the application is all preprocessed offline and stored in a non-relational database. Guide search and off-target scoring is performed using FlashFry ([10]).

## 3 Discussion

Our new tool PAVOOC provides a convenient means to design optimal sgRNAs for KO and genome editing experiments. A machine-learning based scoring system guides the user to select sgRNAs with possibly strong on- and low off-target effects. Through the integration of structural data, PAVOOC is able to display cutting sites on corresponding protein crystal structures such that sgRNAs can be selected which cut in functionally relevant regions. Integration of cancer cell line data ensures that existing genomic alterations are considered during sgRNA selection. The tool was used internally to design a domain-targeting genome-wide sgRNA library.

PAVOOC is hosted on GitHub and is an actively maintained project. As such, it provides an open platform to build and integrate use cases of CRISPR that are not part of the current state. The PEP8 compliant Python code and the react.js-based frontend simplify the entry for developers. The application runs in a Docker environment which makes it easy to host the application on premise.

## Acknowledgements

We thank Robin Winter, Claudia Noack and Barbara Nicke for useful suggestions and discussions throughout the project.

